# MicroRNA-202 prevents precocious spermatogonial differentiation and meiotic initiation during mouse spermatogenesis

**DOI:** 10.1101/2021.06.27.449895

**Authors:** Jian Chen, Chenxu Gao, Xiwen Lin, Yan Ning, Wei He, Chunwei Zheng, Daoqin Zhang, Lin Yan, Binjie Jiang, Yuting Zhao, Md Alim Hossen, Chunsheng Han

## Abstract

Spermatogonial differentiation and meiotic initiation during spermatogenesis are tightly regulated by a number of genes including those coding enzymes for miRNA biogenesis. However, whether and how single miRNAs regulate these processes remain unclear. Here, we report that *miR-202*, a member of the *let-7* family, prevents precocious spermatogonial differentiation and meiotic initiation in spermatogenesis by regulating the timely expression of many genes including those for other key regulators. In *miR-202* knockout (KO) mice, the undifferentiated spermatogonial pool is reduced, ultimately causing agametic seminiferous tubules. SYCP3, STRA8 and DMRT6 are expressed earlier in KO mice than in wild-type (WT) littermates, and *Dmrt6* mRNA is a direct target of *miR-202-5p*. Moreover, the precocious spermatogonial differentiation and meiotic initiation were also observed in KO spermatogonial stem cells when cultured and induced *in vitro*, and could be rescued by the knockdown of *Dmrt6*. Therefore, we have not only shown that *miR-202* is a novel regulator of meiotic initiation but also added a new module to the underlying regulatory network.

**Summary statement:** A single miRNA, *miR-202*, prevents precocious differentiation and meiotic initiation during spermatogenesis. *miR-202*, DMRT6 and STRA8 act together as a novel module in the regulatory network of meiotic initiation.

## Introduction

Spermatogenesis begins with the proliferation and differentiation of spermatogonia and ends with a huge population of spermatozoa. Mouse spermatogonia undergo 8 to 9 consecutive mitotic divisions to generate many subtypes including type A (SG-A), intermediate (SG-I), and type B (SG-B) spermatogonia (de Rooij and Russell, 2000; Griswold, 2016). Consequently, it is estimated one spermatogonial stem cell (SSC) produces about 4096 spermatozoa at the end of spermatogenesis (de Rooij and Russell, 2000). These subtypes can also be roughly divided into undifferentiated (SG_undiff_) and differentiating spermatogonia (SG_diff_). While SG_undiff_ are marked by the expression of PLZF (Buaas et al., 2004; Costoya et al., 2004), SG_diff_ are positive for KIT, a well-known marker for transit-amplifying cells in some other tissues (Besmer et al., 1993). SSCs are a functionally defined subpopulation of SG_undiff_ without specific markers (La et al., 2018; Yoshida, 2019). The transition from SG_undiff_ to SG_diff_ occurs cyclically, resulting in the concurrent presence of several types of cells originated from overlapping cycles to form a fixed number of unique associations (12 in mice) known as seminiferous stages (Russell et al., 1990). Spermatogonial differentiation ends with mitosis-to-meiosis transition, also known as meiotic initiation, which is the hallmark for gametogenesis (de Rooij and Russell, 2000; Griswold, 2016). The disruption of spermatogonial differentiation reduces sperm population (Hobbs et al., 2012; Maezawa et al., 2017). On the other hand, the precocious differentiation/meiosis leads to exhaustion of the spermatogonial pool (Matson et al., 2010).

Spermatogonial development is under the control of both extracellular and intracellular factors. For example, a low level of glial-cell-line-derived neurotrophic factor (GDNF) in GDNF^+/−^ mice was found to result in Sertoli-cell-only seminiferous tubules whereas its overexpression in transgenic mice resulted in an over-proliferation of spermatogonia and testicular tumors (Meng et al., 2000). Retinoic acid (RA) is the key molecule to induce spermatogonial differentiation and meiotic initiation by activating the expression of a large number of genes (Bowles et al., 2006; Koubova et al., 2006; Anderson et al., 2008a; Endo et al., 2015; Wang et al., 2016; Griswold and Hogarth, 2018). The stimulated by retinoic acid gene 8 (*Stra8*) is a direct target and mediator of RA signaling, and plays an essential role in meiotic initiation in both males and females (Oulad-Abdelghani et al., 1996; Anderson et al., 2008a; Griswold and Hogarth, 2018; Ishiguro et al., 2020). STRA8 is also shown to promote spermatogonial differentiation (Endo et al., 2015).

A number of regulators for spermatogonial differentiation and/or meiotic initiation have also been identified, including transcriptional/epigenetic regulating factors such as SOX3 (Raverot et al., 2005), SOHLH1 (Ballow et al., 2006), SOHLH2 (Hao et al., 2008), DMRT1 (Matson et al., 2010), SALL4 (Hobbs et al., 2012), DMRT6 (Zhang et al., 2014), MAX (Suzuki et al., 2016), PRC1 (Maezawa et al., 2017) and MEIOSIN (Ishiguro et al., 2020), RNA binding proteins such as NANOS2 (Suzuki and Saga, 2008), DAZL (Lin et al., 2008), AGO4 (Modzelewski et al., 2012) and BCAS2 (Liu et al., 2017), and ubiquitylation-related protein β-TrCP (Nakagawa et al., 2017). DMRT proteins are transcription factors, and some members have been reported to participate in multiple steps of mammalian spermatogenesis (Zhang and Zarkower, 2017). DMRT1 is required in the establishment and maintenance of SSCs (Zhang et al., 2016), and prevents the precocious activation of the mitosis-meiosis switch (Matson et al., 2010). DMRT6 acts in SG_diff_ to coordinate an orderly transition from the mitotic program to the meiotic program (Zhang et al., 2014).

MiRNAs are fundamental to the developmental, physiologic, and disease processes of metazoans via direct posttranscriptional repression of mRNA targets (Bartel, 2018). In somatic stem/progenitor cells, miRNA function is intricately regulated to promote and stabilize cell fate determination (Shenoy and Blelloch, 2014). Whereas global miRNA loss resulting from KO/mutations of key regulators in miRNA biogenesis induces dramatic phenotypic changes in almost all examined tissues, mice with individual miRNA KOs often lack dramatic phenotypic consequences, implying that miRNAs act in a redundant manner (Park et al., 2012; Shenoy and Blelloch, 2014). MiRNAs are in fact believed to be essential for mammalian spermatogenesis based on the infertile phenotypes of gene KO mice, including KOs of *Dicer*, *Drosha*, and *Dgc8*, which encode key regulators of miRNA biogenesis (Hayashi et al., 2008; Maatouk et al., 2008; Korhonen et al., 2011; Romero et al., 2011; Greenlee et al., 2012; Wu et al., 2012; Zimmermann et al., 2014; Modzelewski et al., 2015; Hilz et al., 2016). However, there have been no reports on the *in vivo* functions of single miRNAs in spermatogenesis based on mouse genetic studies.

The intergenic miRNA gene *miR-202* belongs to the highly conserved *let-7* family (Roush and Slack, 2008) that generates *miR-202-3p* and *miR-202-5p*, which are highly expressed in mouse testes. We previously reported that *miR-202* played an important role in maintaining the stem cell state of cultured mouse spermatogonia (Chen et al., 2017). In the present study, we investigated the *in vivo* function of *miR-202* in the spermatogonial differentiation and meiotic initiation by using KO mice. We found that both spermatogonial differentiation and meiotic initiation occurred precociously upon *miR-202* KO. We also identified that SYCP3, STRA8 and DMRT6 were aberrantly and early expressed in the absence of *miR-202*, and found that DMRT6, whose mRNA was a direct target of *miR-202-5p*, mediated the function of *miR-202*. Our results contribute to our understanding of how a single miRNA safeguards the spermatogonial differentiation and meiotic initiation.

## Results

### *MiR-202* knockout reduces the undifferentiated spermatogonial pool

The *miR-202* KO mice were produced by CRISPR-Cas9 technology, as described in detail in another study (Chen et al., 2021 preprint). No apparent defects were observed in the seminiferous tubules of KO mice at 2 months of age (Fig. S1A), but loss of germ cells became progressively more severe in an age-dependent manner (Fig. 1A and S1A). At 12 months of age, some KO seminiferous tubules were agametic, containing only the somatic Sertoli cells (Fig. 1B and S1B). This phenotype was reminiscent of that of the KO mice of PLZF, a transcriptional factor expressed in SG_undiff_. PLZF is required for the maintenance of the SSC pool, as its KO mice are depleted of germ cells in an age-dependent manner (Buaas et al., 2004; Costoya et al., 2004). Therefore, we examined the expression changes of PLZF in *miR-202* KO mice and found that the PLZF^+^ SG_undiff_ were reduced by 43% (Fig. 1C–E). However, the numbers of differentiating spermatogonia represented by the KIT^+^ cells inside the tubules were similar in the KO and WT mice (Fig. S2A-C), showing that the ability of spermatogonial differentiation was not disturbed in KO mice. These results indicated that *miR-202* KO reduced the pool of SG_undiff_.

**Fig. 1.**
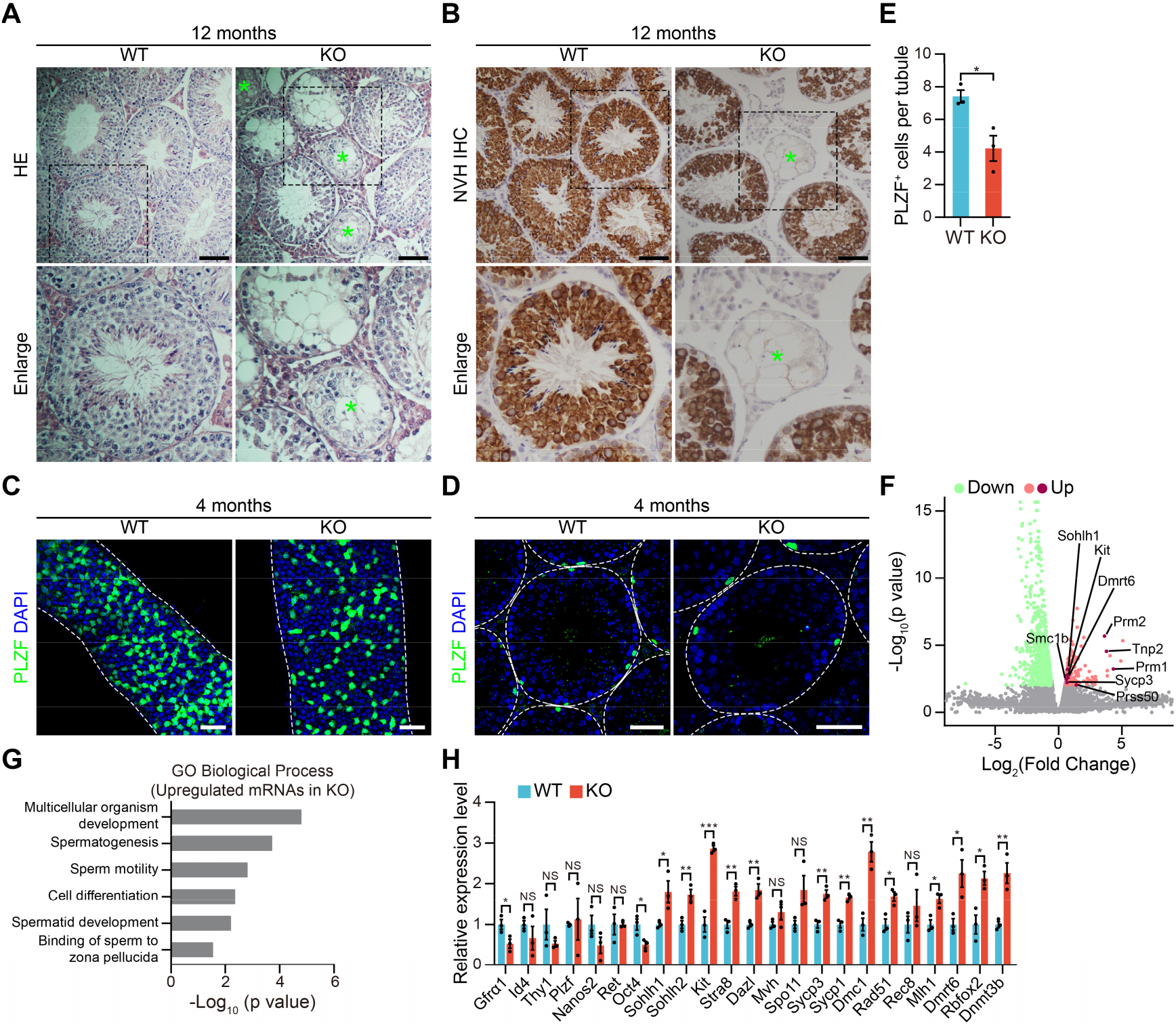
*miR-202* knockout reduces the undifferentiated spermatogonial pool. (**A**) Histological analysis of testis sections from WT and KO mice by H&E staining at 12 months of age. Magnified views are indicated by dashed areas. Green asterisks indicate the agametic tubules. (**B**) Immunohistochemistry of the germ cell marker MVH in testicular sections from mice at 12 months of age. Magnified views are indicated by dashed areas. Green asterisks indicate the agametic tubules. (**C** and **D**) Immunofluorescent staining for PLZF in whole-mount tubules (C) or sections (D) from WT and KO mice at four months of age. Dotted white line, outline of seminiferous tubules or testis tubules. (**E**) Average numbers of PLZF-positive cells per tubule in (D). At least 50 tubules were counted for each mouse. (**F**) Volcano plots showing differentially expressed genes between WT and KO SG-A. Upregulated genes are indicated by red dots and downregulated genes are indicated by green dots. (**G**) Representative Gene Ontology (GO) terms of the biological process categories enriched in upregulated mRNAs in KO SG-A, as identified by RNA-seq. (**H**) qPCR analysis of gene expression in WT and KO SG-A. Data are normalized to WT SG-A. All panels show mean ± SEM, *p < 0.05, **p < 0.01, ***p < 0.001. NS, not significant. Scale bars, 50 μm.

### *MiR-202* knockout induces differentiation- and meiosis-related gene transcription in spermatogonia

To examine whether reduction in KO SG_undiff_ would be caused by premature differentiation/meiosis *in vivo*, we next executed RNA-seq analysis on SG-A (Fig. S3A-B and Table S1), and identified 146 upregulated and 721 downregulated genes resulting from *miR-202* KO (Fig. 1F, Table S1). The upregulated genes were enriched with GO terms including spermatogenesis, sperm motility, cell differentiation, spermatid development and binding of sperm to zona pellucida (Fig. 1G, Table S1). Gene-set enrichment analysis (GSEA) also revealed that genes involved in the GO term of sperm motility were significantly upregulated in KO SG-A (Fig. S3C). Moreover, using qRT-qPCR we validated the downregulation of genes involved in maintaining the undifferentiated spermatogonia (including *Gfra1* and *Oct4*), and the upregulation of genes promoting spermatogonial differentiation and meiotic initiation/progression (such as *Sohlh1*, *Sohlh2*, *Kit*, *Stra8*, *Dazl*, *Spo11*, *Sycp3*, *Sycp1*, *Dmc1*, *Rad51*, and *Mlh1*) in KO SG-A (Fig. 1H) (La et al., 2018). These results suggested that the reduction of SG_undiff_ pool was caused by precocious spermatogonial differentiation and meiotic initiation.

### *MiR-202* knockout results in precocious spermatogonial differentiation and meiotic initiation

We then examined the timing of meiotic initiation by using pre-pubertal mice, in which the first wave of spermatogenesis occurs in a synchronized manner (de Rooij and Russell, 2000). In WT mice at postnatal day 9 (P9), only ~20% of the tubules contained cells of SYCP3^+^, which marked meiotic spermatocytes (Mahadevaiah et al., 2001), whereas in the KO mice, about 72% of tubules contained positive cells (Fig. 2A-B and S4A). SYCP3 and γH2AX co-staining of chromosomal spreads (Mahadevaiah et al., 2001) from mice at P9 showed that the most advanced spermatogenic cells in the WT mice were leptotene spermatocytes, whereas 23% of spermatocytes were zygotene spermatocytes in the KO mice at this time (Fig. 2C-D). These results supported our hypothesis that meiosis was precociously initiated.

**Fig. 2.**
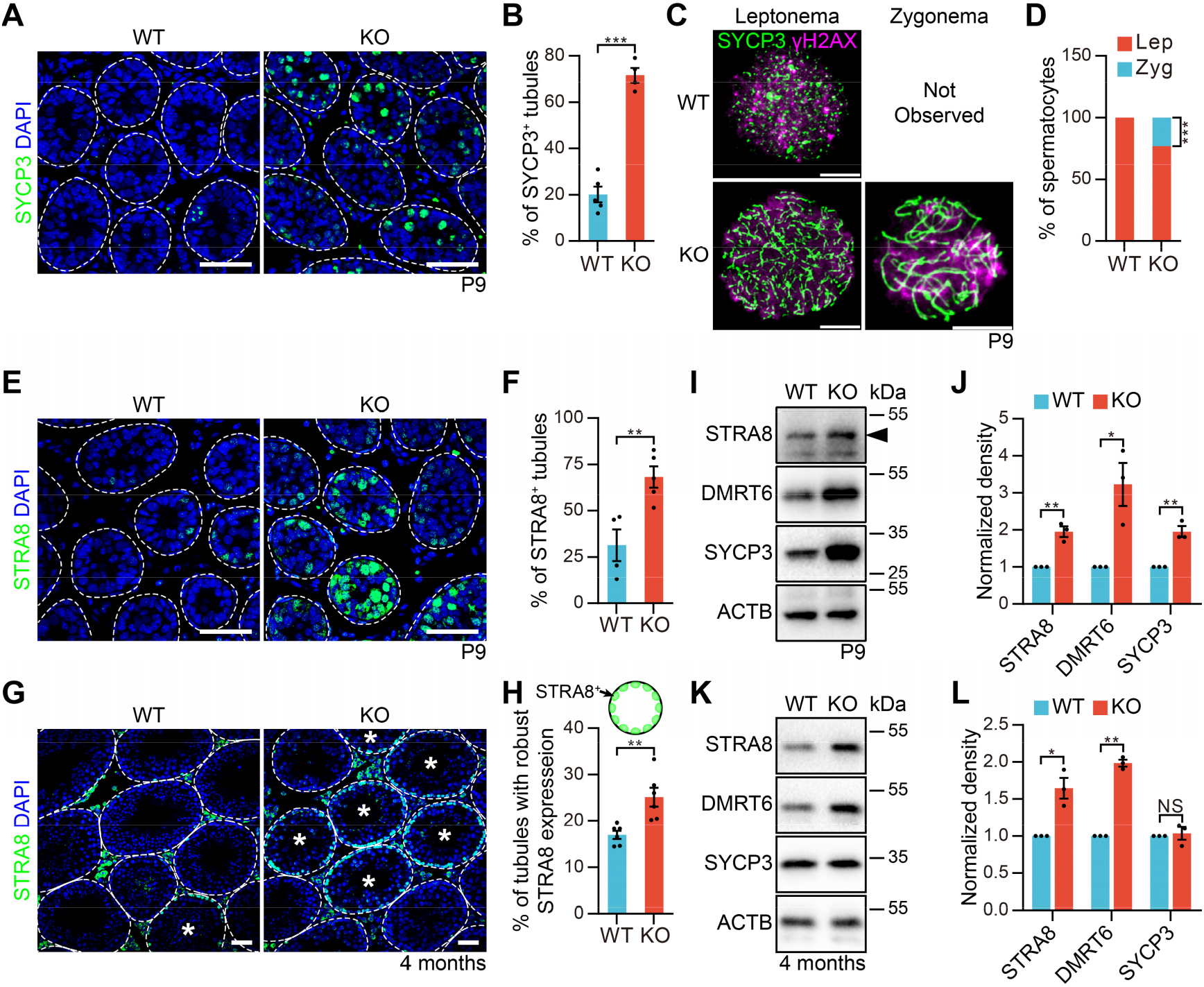
*miR-202* knockout induces precocious meiotic initiation. (**A**) Immunofluorescent staining for SYCP3 in histological sections from mice at P9. Dotted white line, outline of testis tubules. (**B**) Proportion of tubules containing SYCP3^+^ cells in (A). At least 50 tubules were counted for each mouse. (**C**) Immunofluorescent staining for SYCP3 and γH2AX in nuclear spreads of spermatocytes from mice at P9. (**D**) Proportion of leptonema and zygonema in (C). n = 3. (**E** and **G**) Immunofluorescent staining for STRA8 in sections from mice at P9 and 4 months of age (G). White asterisks indicate tubules with robust STRA8 expression in (G). Dotted white line, outline of testis tubules. (**F** and **H**) Proportion of tubules containing STRA8-positive cells at P9 (F) and tubules with robust STRA8 expression at 4 months of age (H). The upper panel in (H) is the schematic representation for robust STRA8 expression. At least 50 tubules were counted for each mouse. (**I** and **K**) Western blotting analyses of WT and KO testes from mice at P9 (I) and 4 months of age (K), n = 3. (**J** and **L**) The normalized densities in (I) and (K), respectively. All panels show mean ± SEM, *p < 0.05, **p < 0.01, ***p < 0.001. NS, not significant. Scale bars, 50 μm.

To further verify the observations in KOs, we examined the expression pattern of STRA8 as it is a key factor promoting spermatogonial differentiation and meiotic initiation (Anderson et al., 2008b; Mark et al., 2008; Endo et al., 2015; Ishiguro et al., 2020). Compared with the WT mice, testis sections from KO mice at P9 contained more STRA8^+^ tubules (~2.2-fold increase) (Fig. 2E-F and S4B). The tubules with robust STRA8 expression in adult KO mice were also significantly increased compared with the WT littermates (25% vs. 17%; Fig. 2G-H). The upregulation of STRA8 was confirmed through western blot analyses, which displayed 2- and 1.6- fold increases in P9 and adult testes, respectively (Fig. 2I-L and S9A-B).

We next wondered how the increased STRA8 level arose by co-immunostaining for STRA8 and PLZF. STRA8 is expressed at low levels in spermatogonia and at greatly elevated levels in preleptotene spermatocytes (pre-lep) where mitosis-meiosis transition occurs (Zhou et al., 2008). STRA8 expression overlaps with PLZF in stages VII-VIII, and is limited to PLZF-low and -negative spermatogonia thereafter in stages IX-X (Endo et al., 2015). We found that more PLZF^+^ cells became STRA8^+^ cells in KO than in WT at P9 and 4 months of age (Fig. 3A-D). Moreover, in adult mice, STRA8 expression also overlapped with PLZF in stages I-VI in KOs, showing the widespread and early presence of STRA8 independent of the seminiferous epithelial cycle (Fig. 3C, E). Taken together, the loss of *miR-202* in SG_undiff_ caused precocious spermatogonial differentiation and meiotic initiation.

**Fig. 3.**
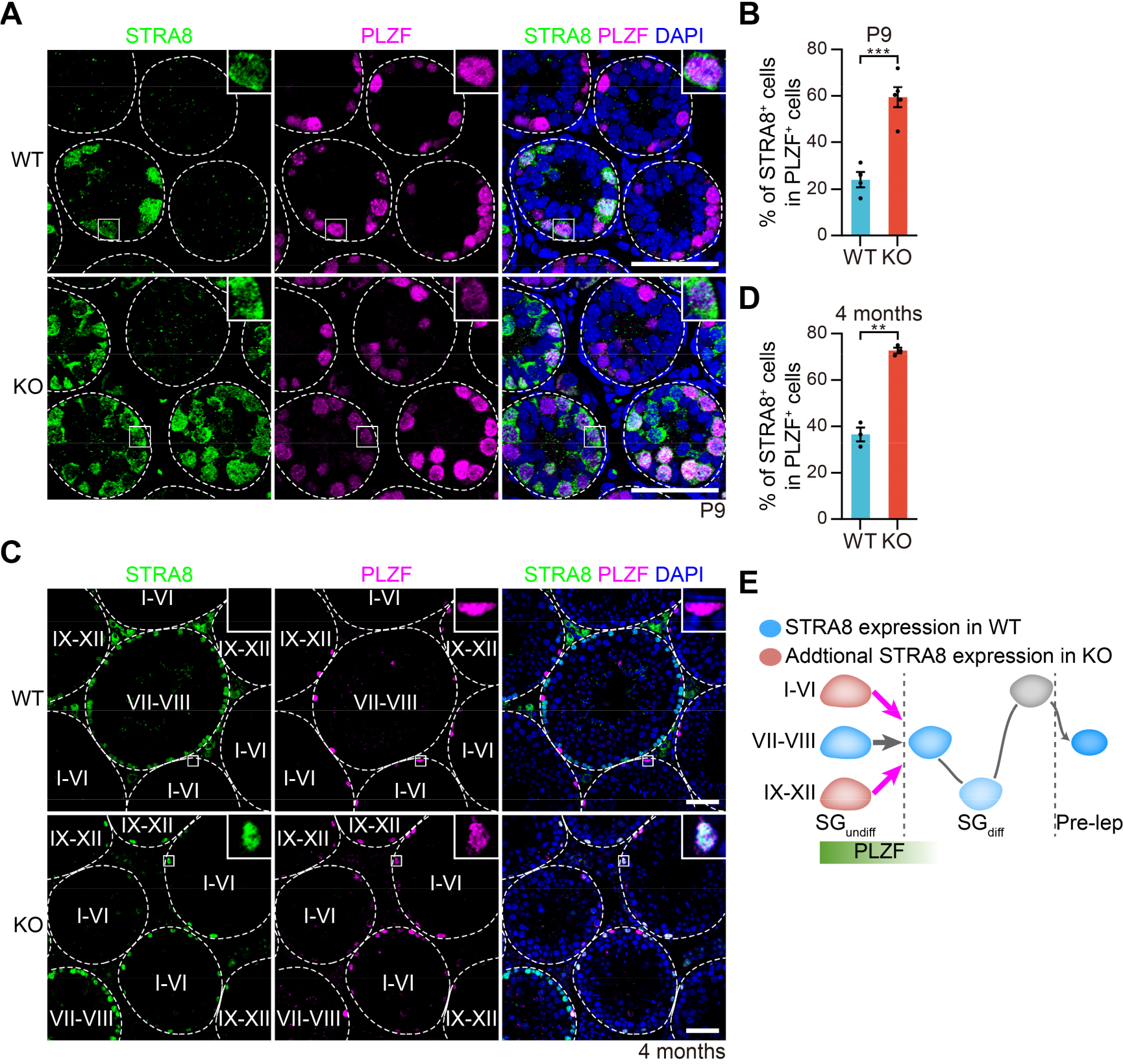
*miR-202* knockout induces early and widespread presence of STRA8 in undifferentiated spermatogonia. (**A** and **C**) Immunofluorescent staining for STRA8 and PLZF in sections from mice at P9 (A) and 4 months of age (C). Insets show high-magnification images of boxed regions. The seminiferous tubules in adult sections (C) were classified into three types based on their seminiferous stages. Dotted white line, outline of testis tubules. (**B** and **D**) The percentage of STRA8^+^ cells in PLZF^+^ undifferentiated spermatogonia in (A) and (C), respectively. At least 50 tubules were counted for each mouse. (**E**) Diagram of STRA8 expression in SG_undiff_, SG_diff_ and preleptonema, in WT (blue) and KO (blue + pink) mice. Magenta arrows indicate the uncontrolled transition from undifferentiated to differentiating spermatogonia in KO. Diagram is based on observations in (C). All panels show mean ± SEM, **p < 0.01, ***p < 0.001. NS, not significant. Scale bars, 50 μm.

### DMRT6 expression is pre-activated in the absence of *miR-202*

We subsequently wondered how *miR-202* mediated the precocious meiotic initiation. As *Stra8* mRNA is not a predicted *miR-202* target (Paraskevopoulou et al., 2013; Agarwal et al., 2015), we focused on *Dmrt6* because: 1) *Dmrt6* plays a crucial role in the mitosis/meiosis switch (Zhang et al., 2014); 2) its 3’ UTR contains an *miR-202-5p*-binding site (Paraskevopoulou et al., 2013); and 3) its expression was upregulated in *miR-202* KO SG-A (Fig. 1F, H). Similar to what was observed for STRA8, the KO testes contained more DMRT6^+^ tubules (~a 2.6-fold increase) relative to the WT tubules at P9 (Fig. 4A-B and S4C). The adult KO mice also contained more DMRT6^+^ tubules than the WT (75% vs. 68%; Fig. 4C-D), and the upregulation of DMRT6 was also confirmed through western blot analyses in P9 and adult testes (Fig. 2I-L and S9A-B). M*iR-202-5p* was one of the five most abundant miRNAs in SG-A (~7% of total reads, Fig. 4E), according to our RNA sequencing data (Chen et al., 2017). Dual luciferase assay revealed that the 3’ UTR of *Dmrt6* mRNA that contained the predicted target site was actually targeted by the *miR-202-5p* mimic, and as a negative control, the mutated target sequence was not targeted by *miR-202-5p*, further indicating that *Dmrt6* was a direct target of *miR-202-5p* (Fig. 4F). As *miR-202* mutants mostly phenocopied mice lacking DMRT1, which was reported to prevent the precocious mitosis/meiosis switch and was expressed in SG and Sertoli cells (Matson et al., 2010), we examined the DMRT1 expression pattern in our KO mice, and found no obvious differences between WT and KO mice (Fig. S5A-B).

**Fig. 4.**
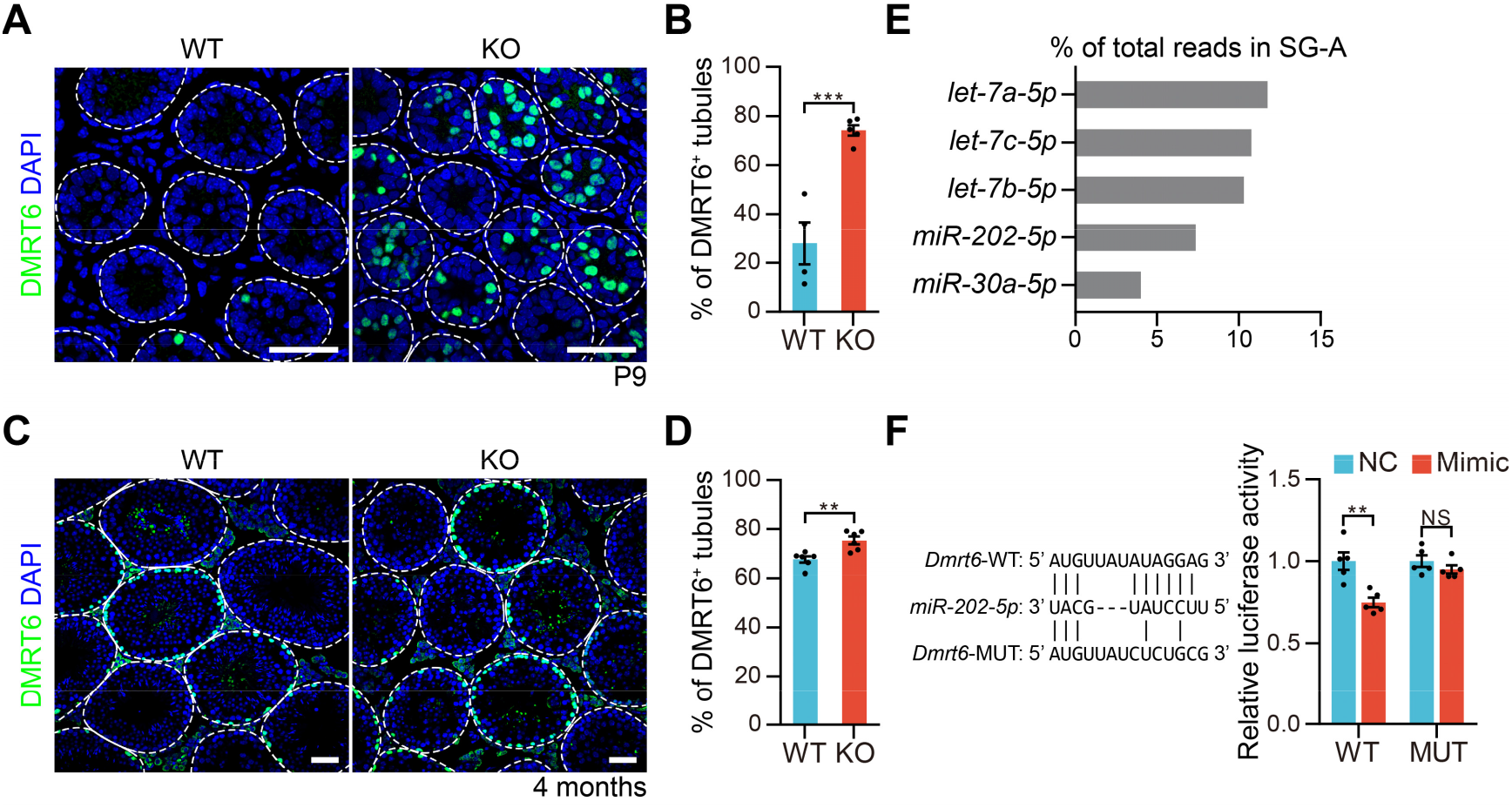
*miR-202* knockout leads to DMRT6 pre-activation. (**A** and **C**) Immunofluorescent staining for DMRT6 in sections from mice at P9 (A) and 4 months of age (C). Dotted white line, outline of testis tubules. (**B** and **D**) Proportion of tubules containing DMRT6-positive cells at P9 (B) and 4 months of age (D). At least 50 tubules were counted for each mouse. (**E**) Percentage of total reads for the five most abundant miRNAs in SG-A. Note that *miR-202-5p* occupies ~7% of total reads. Validation of *Dmrt6* mRNA as a direct target of *miR-202-5p* by dual luciferase assay. Data are normalized to the scrambled negative control (NC) mimic. Left panel, the sequence of *miR-202-5p*, and predicted miRNA regulatory elements at the 3’ UTR of *Dmrt6* mRNA and mutated 3’ UTR sequence. All panels show mean ± SEM, **p < 0.01, ***p < 0.001. NS, not significant. Scale bars, 50 μm.

To distinguish whether DMRT6 was actually pre-activated in SG_undiff_, we compared the expression pattern of DMRT6 with that of PLZF in P9 and adult mice (Fig. 5A-B). DMRT6 was only expressed in SG_diff_, showing no overlap with the SG_undiff_ marker PLZF (Zhang et al., 2014), and we indeed observed that DMRT6 and PLZF did not co-localize in WT mice (Fig. 5A-B). However, in KOs the DMRT6 and PLZF partially co-localized, indicating that DMRT6 started to be expressed in cells as early as SG_undiff_ (Fig. 5A-B). We also wished to ascertain when DMRT6 would end its expression using co-immunostaining for both DMRT6 and SYCP3 (Fig. S6A-B). DMRT6 and SYCP3 staining was mutually exclusive in juvenile and adult WT mice, whereas a small number of cells were double-positive in KOs (Fig. S6A-B). In KO adult mice, DMRT6 expression overlapped with SYCP3 in stages VII-VIII, specifically in pre-lep (Fig. S6B). These results indicated that in the absence of *miR-202*, DMRT6 was pre-activated in SG_undiff_, and continued to be expressed in SYCP3^+^ preleptonema. Such an extended expression window may contribute to premature spermatogonial differentiation and meiotic initiation.

**Fig. 5.**
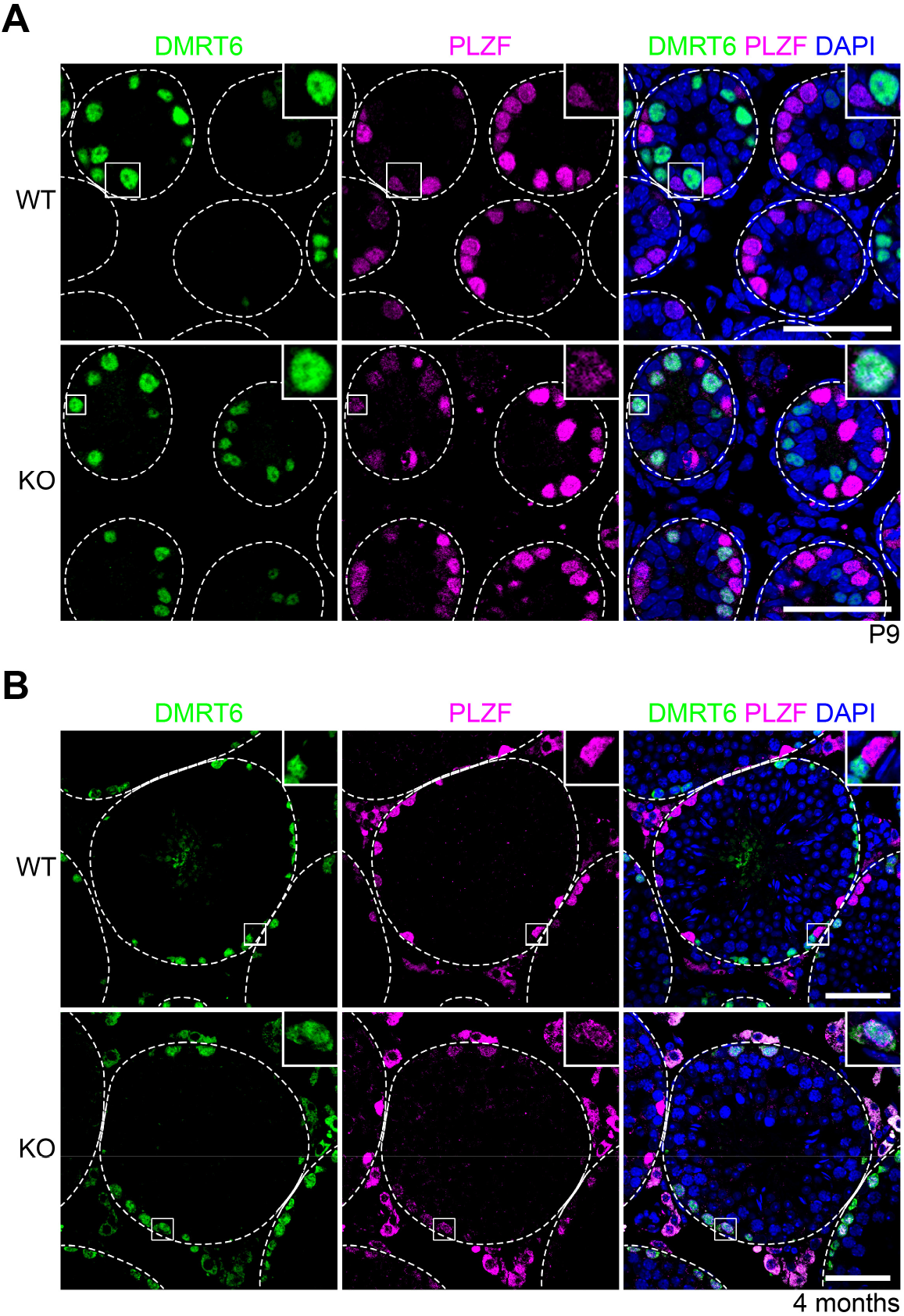
*miR-202* knockout induces precocious expression of DMRT6 as early as in SG_undiff_. (**A** and **B**) Immunofluorescent staining of sections from mice at P9 (A) and 4 months of age (B) for DMRT6 and PLZF. Insets show high magnification images of boxed regions. Dotted white line, outline of testis tubules. Scale bars, 50 μm.

### Loss of *miR-202* primes cultured undifferentiated spermatogonia for spermatogonial differentiation and meiotic initiation

To further test our hypothesis that the *miR-202*/DMRT6 axis regulates the initiation of meiosis, we used SSC lines established in in another study (Chen et al., 2021 preprint). The establishment rate of KOs was significantly lower than that of the WT mice (3/7 vs. 11/11; Chi-squared test, p<0.005). The cells of these SSCs were more appropriately named SG_undiff_ because only a small fraction were bona-fide SSCs based upon transplantation assays (Kanatsu-Shinohara et al., 2003; Kubota et al., 2004). Using these cell lines, we first assessed proliferation, and found that KO SG_undiff_ were more mitotically active than WT SG_undiff_ based on cell counting and BrdU-incorporation assays (Fig. 6A, S7A-B). However, KO SG_undiff_ also exhibited a higher apoptotic rate, as revealed by TUNEL assays (Fig. 6B, S7C). When we compared the stem cell activity of the WT and KO SG_undiff_ by transplantation assays (Fig. 6C), we found that KO SG_undiff_ exhibited 25% lower stem cell activity than WT (Fig. 6D), consistent with our previous report (Chen et al., 2017). These results indicated that *miR-202* facilitated the maintenance of a more stem cell-like state of SG_undiff_ that possessed lower mitotic and apoptotic rates and higher stem cell activity.

**Fig. 6.**
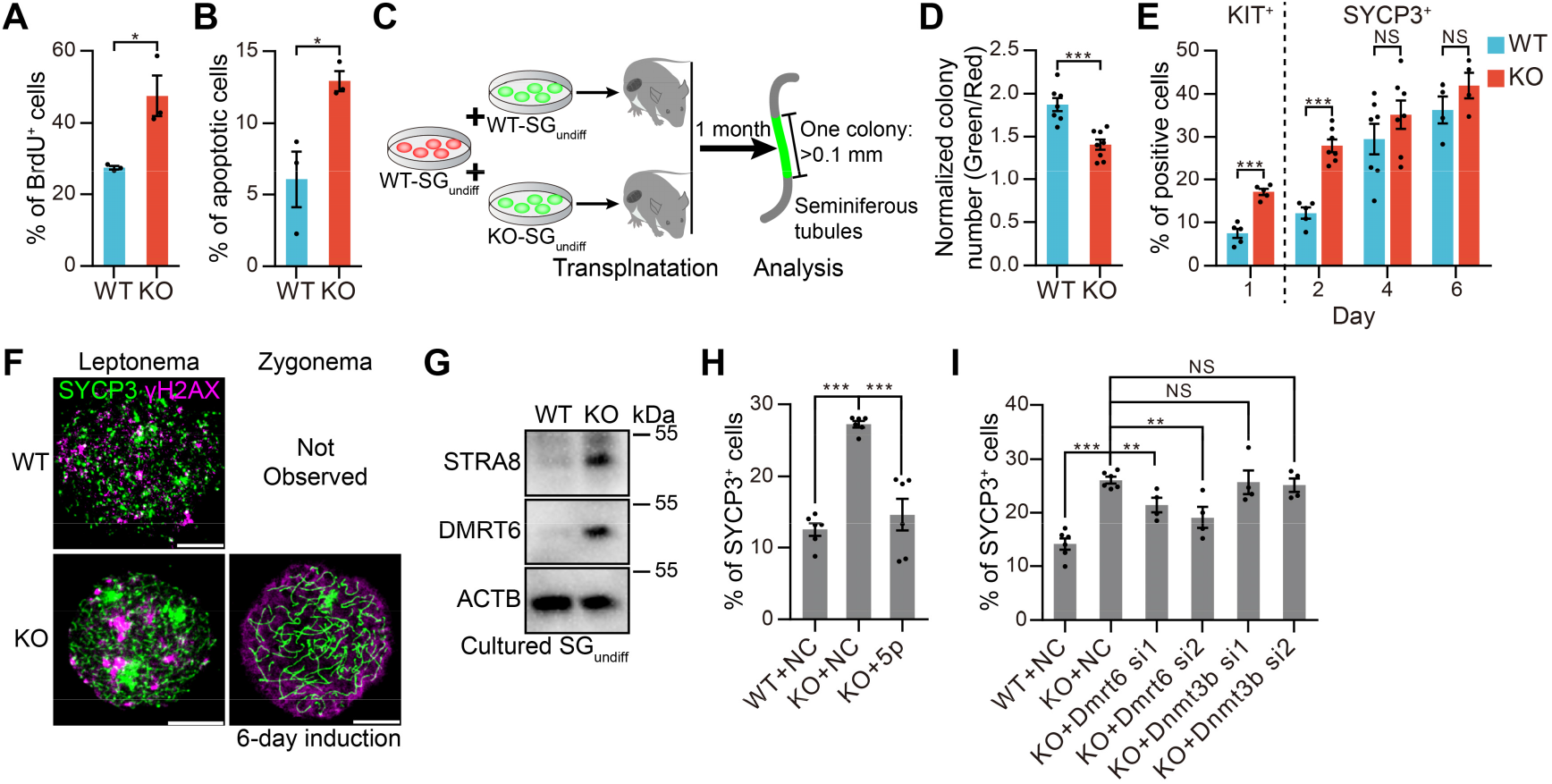
*miR-202* knockout primes undifferentiated spermatogonia for differentiation and meiotic initiation. (**A** and **B**) Proportion of BrdU^+^ (A) and apoptotic (B) cultured SG_undiff_. At least 400 cells for each biological replicate were counted. (**C**) Scheme of transplantation assay. EGFP-labelled WT or KO cultured SG_undiff_ were pooled with mRuby2-labelled WT SG_undiff_, and the cells were transplanted into recipient mice. One month later, colony number was counted. (**D**) Quantitative comparison of stem cell content of WT and KO SG_undiff_ by transplantation assay in (C). Note that mRuby2-labelled SG_undiff_ were used as internal controls. (**E**) Proportion statistics of KIT- and SYCP3-positive cells in WT and KO SG_undiff_ by flow cytometric analyses after meiotic induction on Sertoli cells for the indicated time. (**F**) Nuclear spread staining of WT and KO SG_undiff_ for SYCP3 and γH2AX after induction on Sertoli cells for 6 days. n = 3. (**G**) Western blotting analyses in cultured SG_undiff_, n = 3. (**H** and **I**) Proportion statistics of SYCP3-positive cells in WT and KO SG_undiff_ by flow cytometric analyses after pretreatment with mimics (H) or siRNAs (I) for 10 hours and induction on Sertoli cells for two days. All panels show mean ± SEM, *p < 0.05, **p < 0.01, ***p < 0.001. NS, not significant. Scale bars, 10 μm.

We next used an *in vitro* meiosis-initiation model to further investigate the mechanisms of *miR-202* function. Since SG_undiff_ cultures can be induced to initiate meiosis when they are plated on a neonatal Sertoli feeder layer (Wang et al., 2016), We monitored the differentiation and meiotic initiation by examining the expression of KIT and SYCP3 (Fig. 6E, S7D). One day after induction, we found that 18% of the KO cells started to expresses KIT, whereas only 7% WT cells did so. On day 2 of induction, more KO cells than WT cells began to express SYCP3 (28% vs. 12%; Fig. 6E, S7D). We further conducted co-staining of SYCP3 and γH2AX on chromosome spreads to examine the types of WT and KO SYCP3^+^ cells that were generated 6 days after induction, and found that the most advanced spermatogenic cells in KO were zygotene spermatocytes, in contrast to leptotene spermatocytes in WT (Fig. 6F). These *in vitro* results again indicated that KO SG_undiff_ can be induced to initiate differentiation and meiosis more readily, and that *miR-202* deletion predisposed SG_undiff_ to a differentiation- and meiosis-primed state.

### *MiR-202* function in meiotic initiation is mediated by *Dmrt6*

When we examined whether STRA8 and DMRT6 were pre-activated in KO SG_undiff_ which were PLZF^+^, we indeed detected both STRA8 and DMRT6 in KOs but not in WT, consistent with our previous results (Fig. 6G, S7E). We further tested whether the precocious meiotic initiation of *miR-202* KO SG_undiff_ *in vitro* could be rescued by *miR-202-5p* mimic or by the knockdown of *Dmrt6* mRNA. We selected *Dnmt3b* (which is involved in DNA methylation) as a potential negative control based on that: 1) *Dnmt3b* mRNA was upregulated in KO SG-A (Fig. 1H, Table S1); 2) *Dnmt3b* was a direct target of *miR-202-3p* using dual luciferase assays (Fig. S8A) (Agarwal et al., 2015); and 3) *Dnmt3b* conditional KO in germ cells manifested no phenotypic abnormality in spermatogenesis (Kaneda et al., 2004). SG_undiff_ cultures were pretreated with *miR-202-5p* mimic or with siRNAs targeting *Dmrt6* and *Dnmt3b* (Fig. S8B) before induction was conducted. On day 2 of induction, about 27% of the KO cells pretreated with control mimic and siRNA were SYCP3^+^, which was similar to the percentage without pretreatment (Fig. 6E, H and I); this percentage was reduced to ~15% by pretreatment with the *miR-202-5p* mimic, which approximated the value observed in the WT cells pretreated with a scrambled control (Fig. 6H). Moreover, knockdown of *Dmrt6* also reduced the percentages of SYCP3^+^ cells significantly, while knockdown of *Dnmt3b* did not (Fig. 6I). These *in vitro* results thereby indicated that the function of *miR-202* in inhibiting precocious meiotic initiation was mediated by its downstream target, *Dmrt6*.

## Discussion

Spermatogonial differentiation and meiotic initiation must be precisely regulated to ensure both the productive generation of spermatozoa and the maintenance of the stem cell pool to support the life-long spermatogenesis (Griswold, 2016; Griswold and Hogarth, 2018). However, only a small number of regulators for these two important processes have been identified. In the present study, we found that the KO of *miR-202* resulted in precocious spermatogonial differentiation, meiotic initiation, and a reduced SG_undiff_ pool. We also demonstrated that loss of *miR-202* results in pre-activation of STRA8 and DMRT6, two key factors promoting differentiation and meiosis (Oulad-Abdelghani et al., 1996; Anderson et al., 2008a; Zhang et al., 2014; Endo et al., 2015; Griswold and Hogarth, 2018; Ishiguro et al., 2020), in SG_undiff_, and found that *Dmrt6* mRNA was directly targeted by *miR-202-5p*. Therefore, a module composed of *miR-202*, DMRT6 and STRA8 has been identified in the context of the spermatogonial fate-decision regulatory network (Fig. 7A).

**Fig. 7.**
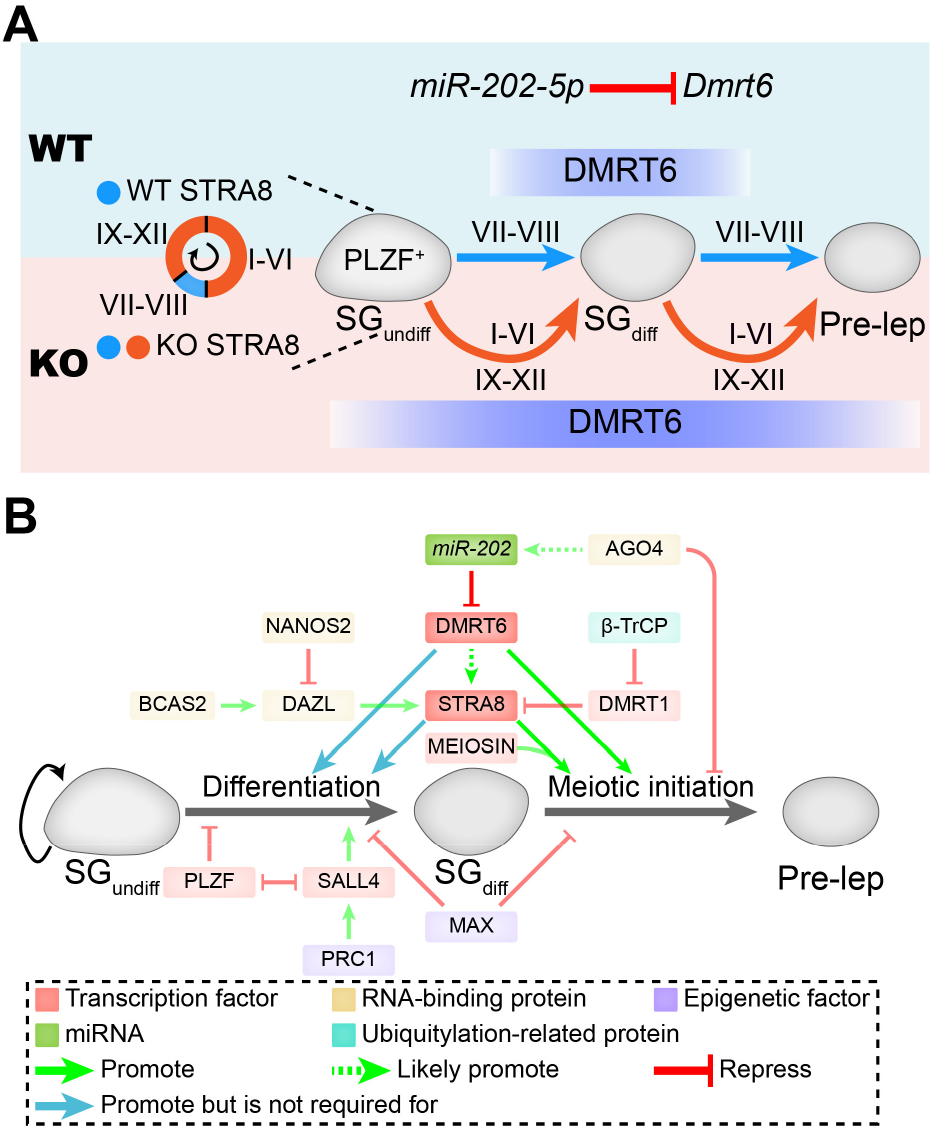
*miR-202*, DMRT6 and STRA8 act together as a novel module in the regulatory network of spermatogonial differentiation and meiotic initiation. (**A**) *MiR-202* prevents precocious spermatogonial differentiation and meiotic initiation. In WT mice, *miR-202* directly targets *Dmrt6* mRNA and restricts the expression window of DMRT6 in SG_diff_, to coordinate an orderly transition from the mitotic program to the meiotic program. The transitions of spermatogonial differentiation and meiotic initiation cooccur in stages VII-VIII, with expression of STRA8. Upon *miR-202* KO, DMRT6 is precociously expressed as early as in SG_undiff_, causing precocious spermatogonial differentiation and meiotic initiation, and a reduced SG_undiff_ pool, with the presence of STRA8 in all stages in SG_undiff_. (**B**) The regulatory network composed of *miR-202*, DMRT6, STRA8 and other regulators orchestrates spermatogonial differentiation and meiotic initiation.

The precocious initiation of spermatogonial differentiation and meiotic initiation in *miR-202* KO mice are supported by multiple lines of evidences including the premature expression of SYCP3, STRA8, DMRT6, and upregulation of a large number of meiotic and post-meiotic genes. In WT mice, the expression of STRA8 in SG_undiff_ and pre-lep is restricted in stages VII-VIII of the seminiferous cycle (Endo et al., 2015) while in KO mice, the STRA8 expression was observed in all stages. Moreover, more SG_undiff_ become STRA8^+^ in the absence of *miR-202* (Fig. 3A-D). DMRT6 is normally expressed in SG_diff_ that are negative for PLZF expression (Zhang et al., 2014). However, in the absence of *miR-202*, the expression of DMRT6 was not only augmented in but also detected as early as in PLZF^+^ SG_undiff_ (Fig. 4A-D, 5A-B). These results suggest that the precise control of spermatogonial differentiation and meiotic initiation was disrupted upon *miR-202* KO (Fig. 7A).

The regulators that govern the spermatogonial differentiation and/or meiosis identified so far include transcription factors, epigenetic regulators, RNA-binding proteins, and proteins involved in RA pathways. Polycomb repressive complex 1 (PRC1) promotes spermatogonial differentiation by timely activating the expression of germline genes as the gene KO mice of *Rnf2*, which encodes the catalytic subunit of PRC1, causes severe defects in spermatogonial differentiation (Maezawa et al., 2017). The effect of PRC1 is mediated by SALL4, a transcription factor promoting spermatogonial differentiation by sequestering PLZF from DNA binding (Hobbs et al., 2012; Maezawa et al., 2017). In contrast, MAX, one component of an atypical PRC1 complex (PRC1.6), prevents spermatogonial differentiation and meiotic onset as indicated by the meiosis-like cytological changes induced by its knockdown in cultured germline stem cells (Maeda et al., 2013; Suzuki et al., 2016). Additional meiotic initiation activators include DAZL and MEIOSIN, and suppressors include NANOS2, AGO4 and DMRT1. DAZL is a key regulator to facilitate the meiotic initiation (Lin et al., 2008) and activates the expression of *Stra8* by targeting to the 3’ UTRs of *Stra8* mRNAs (Li et al., 2019). Interestingly, the expression of *Dazl* is repressed by NANOS2, which suppresses meiotic initiation, and activated by BCAS2, which promotes the process. Both NANOS2 and BCAS2 are RNA-binding proteins (Suzuki and Saga, 2008; Kato et al., 2016; Liu et al., 2017). AGO4, a small RNA binding protein, prevents precocious meiotic initiation, and its deletion results in loss of many miRNAs, including *miR-202* (Modzelewski et al., 2012). It is likely that the function of AGO4 in meiotic initiation is at least partially mediated by *miR-202*. MEIOSIN interacts with STRA8 to activate meiotic initiation (Ishiguro et al., 2020). DMRT1 inhibits meiosis in SG_undiff_ by limiting RA-dependent transcription, and by specifically blocking *Stra8* transcription at pre-lep (Matson et al., 2010). β-TrCP, a component of an E3 ubiquitin ligase complex, targets DMRT1 for degradation and thereby activates meiotic initiation (Nakagawa et al., 2017).

M*iR-202* KO mice phenocopy the mutants of DMRT1, NANOS2, MAX, and AGO4 in that premature meiotic initiation is observed. However, some differences occur among them. For example, spermatogonial differentiation in DMRT1 KO mice is truncated resulting in SG_diff_ depletion (Matson et al., 2010), while it does not occur in *miR-202* mutants as shown by the unaffected KIT^+^ SG_diff_ (Fig. S3A-C). NANOS2 suppresses meiosis and activates a male-specific genetic program in gonocytes, indicating that NANOS2 plays a key role earlier than *miR-202*. MAX prevents differentiation/meiosis via interaction with distinct cofactors in cultured cells, and its *in vivo* function warrants further investigation (Maeda et al., 2013; Suzuki et al., 2016).

Based on our findings and published studies, it can be seen that many regulators cooperate for the fate decision of spermatogonial differentiation and meiotic initiation, and *miR-202* and its direct target *Dmrt6* as well as *Stra8*, of which the expression level and timing are also regulated by *miR-202*, act together as a novel module in this expanding regulatory network which is summarized in Fig. 7B. One interesting research direction in the future is to elucidate how the expression and action of *miR-202* are regulated by known or novel players in the network.

## Materials and methods

### Mice

All of the animals used in this study were approved by the Animal Ethics Committee of the Institute of Zoology at the Chinese Academy of Science. All of the procedures were conducted in accordance with institutional guidelines. Animals were specific-pathogen free (SPF). All mice had access to food and water ad libitum, were maintained on a 12:12 hours light-dark artificial lighting cycle, with lights off at 19:00, and were housed in cages at a temperature of 22-24°C.

The *miR-202* KO mice were produced as described in detail in another study (Chen et al., 2021 preprint). All of the mice were maintained on a C57BL/6J;ICR mixed background. The deletion of the *miR-202* cassette was verified by genomic PCR (Primers are listed in Table S2).

### Histological analyses

Testes dissected from the wildtype and KO mice immediately after euthanasia were fixed in 4% paraformaldehyde or Bouin's Solution for up to 24 hours, dehydrated using gradient concentration of ethanol, treated with xylene, and then embedded in paraffin. Five-micrometer-thick sections were cut and mounted on glass slides. After deparaffinization in xylene and re-hydration in gradient concentration of ethanol, the Bouin-fixed testis cross-sections were used for Hemotoxylin and Eosin (HE) staining, and PFA-fixed sections were used for immunohistochemistry (IHC), immunofluorescence (IF) analyses, and TUNEL assays.

### Antibodies

All antibodies used in this study are listed in Table S3. We obtained the anti-DMRT6 and anti-DMRT1 antibodies from Prof. David Zarkower in University of Minnesota (Matson et al., 2010; Zhang et al., 2014; Zhang et al., 2016), as generous gifts.

### Section immunostaining

Deparaffinized sections were boiled for 15 min in a sodium citrate buffer for antigen retrieval. Then, the slides were incubated with primary antibodies overnight at 4°C and then incubated with secondary antibodies for two hours at room temperature. For IF, signals were visualized by conjugating fluorophore with the secondary antibodies. For IHC, the sections were stained with a horseradish peroxidase (HRP)-conjugated secondary antibody.

### Whole-mount immunostaining

The mouse testes were dissected to remove the tunica albuginea, and seminiferous tubules were untangled. Tubules were fixed in 4% PFA overnight at 4°C, permeabilized with 0.1% Triton X-100 in PBS for four hours at room temperature, and blocked with 5% BSA in PBS for two hours at room temperature. The tubules were incubated with primary antibodies overnight at 4°C and subsequently with fluorophore-conjugated secondary antibodies for four hours at room temperature. The nuclei were counterstained with DAPI.

### Spermatocyte chromosome spreads

Chromosome spreads of the testicular samples were performed using the drying-down technique previously described by Peters *et al.* (Peters et al., 1997). Briefly, the testes were dissected, and the seminiferous tubules were washed in PBS. Then, tubules were placed in a hypotonic extraction buffer for 30–60 min. Subsequently, the tubules were torn to pieces in 0.1 M sucrose (pH 8.2) on a clean glass slide and were pipetted repeatedly to make a suspension. The cell suspensions were then dropped on slides containing 1% PFA and 0.15% Triton X-100 (pH 9.2). The slides were dried for at least two hours in a closed box with high humidity. Finally, the slides were washed twice with 0.4% Photo-Flo 200 (Kodak) and dried at room temperature. The dried slides were stored at −20°C for the immunofluorescent staining. Immunolabeled chromosome spread nuclei were imaged on confocal laser scanning microscopes (Leica or Carl Zeiss) using 63× oil-immersion objective.

### Dual luciferase assay

We used TargetScan or microT-CDS to predict mRNA target sites of *miR-202* (Paraskevopoulou et al., 2013; Agarwal et al., 2015). The predicted 3’ UTRs for candidate genes were amplified from mouse testis cDNA (Primers are listed in Table S2) and inserted into the pMIR-REPORT Luciferase vector.

Dual luciferase assays were performed following our previous report (Chen et al., 2017). Briefly, the 293FT cells were plated in 96-well plate. About 24 hours later, the *miR-202* or scrambled NC mimics of 100 nM were first transfected using the Lipofectamine RNAiMAX Reagent (Invitrogen). Ten hours later, 50 ng of the Firefly Luciferase report plasmids (pMIR-REPORT) and 5 ng of the Renilla Luciferase internal control plasmid (pRT-TK) were co-transfected by the Lipofectamine 2000 Reagent (Invitrogen). Forty-eight hours after plasmid transfection, luciferase activity was examined using the Dual-Luciferase Report Assay System (Promega) on the Synergy4 (Bio-Tek) platform. Data were first normalized to the empty the vector and then to the NC mimic.

### Isolation of type A spermatogonia (SG-A)

Mice at 6-7 dpp were used for SG-A isolation through the STAPUT method following our previous report (Gan et al., 2013). Mice were sacrificed, and then the testes were removed and decapsulated. The seminiferous tubules were cut into small pieces. The pieces were incubated in DPBS containing Collagenase I (Gibco) and then in Trypsin (Gibco) containing DNase I. The dispersed cells were filtered through a 40-μm cell strainer (BD Falcon). After filtered, the cells were resuspended in 0.5 ml of DMEM containing 0.5% BSA. Cells were bottom-loaded into a 10-ml syringe, and this was followed by 10 ml of a 2-4% BSA gradient in DMEM. After two hours of velocity sedimentation at unit gravity, the cell fractions (0.5 ml/fraction) were collected from the bottom of the syringe at a rate of about 1 ml/min. The cell type and purity in each fraction were assessed using a light microscope based on their diameters and morphological characteristics. Only fractions with a purity ≥ 85% were pooled together (Fig. S3A-B).

### qRT-PCR

Primers used in qRT-PCR are listed in Table S2. Total RNAs were isolated using Trizol regent (Invitrogen). Total RNAs were reverse-transcribed using High-Capacity cDNA Reverse Transcription Kit (Applied Biosystems) and the reactions were performed using Hieff qPCR SYBR Green Master Mix (Yeasen) in a Roche LightCycler 480 Real-Time PCR system. The data were then analyzed using the comparative Ct method (ΔCt), with β-actin RNA used as the internal control (Gan et al., 2013).

### Cell cultures

All cells were maintained at 37°C under 5% CO2 and tested negative for mycoplasma contamination. Establishment and maintenance of SSCs (SG_undiff_ in this paper) and Sertoli cells was described in detail in another study (Chen et al., 2021 preprint). The Sertoli cells were treated with mitomycin C and used for induction of meiotic initiation. The 293FT cell lines were maintained in DMEM medium supplemented with 10% FBS.

### Western blotting

Cell lysates from mouse testes were prepared by homogenizing small pieces of organs with glass homogenizers in an RIPA buffer (Beyotime) supplemented with Protease Inhibitor Cocktail (Sigma-Aldrich). Cultured SG_undiff_ were directly lysed in RIPA buffer supplemented with Protease Inhibitor Cocktail. Lysates were then centrifuged at 20,000 g for 10 minutes at 4°C, and supernatants were used for Western blot analyses. Briefly, lysates were run in SDS-PAGE gel and transferred to PVDF membranes. The blots were blocked with 5% BSA for 2 hours, incubated with primary antibodies overnight at 4°C and then incubated with HRP-conjugated secondary antibodies at room temperature for 2 hours. The proteins were detected using SuperSignal West Pico PLUS Chemiluminescent Substrate (Thermo Fisher) on the ChemiDoc XRS^+^ system (Bio-Rad). The band densities were analyzed by ImageJ.

### BrdU incorporation assay

The SG_undiff_ were seeded on Laminin-coated glass in a plate and were treated with 10 μg/ml of BrdU for 12 hours. Then, the cells were fixed with 70% ice-cold ethanol, denatured with 2 M of HCl and neutralized with 0.1 M of sodium borate (pH 8.5). Afterward, the cells were analyzed using a standard procedure similar to that used for immunofluorescence.

### Transplantation of SG_undiff_

Transplantation of the SG_undiff_ into recipient testes was performed as described previously (Kubota et al., 2004; Wang et al., 2015). Briefly, and 5×10^4^ internal control SG_undiff_ labelled by mRuby2 together with 10^5^ WT or KO SG_undiff_ labelled by EGFP were mixed and transplanted into each testis of busulfan-pretreated recipient mice. One month later, the colony number was counted and analyzed.

### Meiotic initiation induction of SG_undiff_

SGundiff were digested with Accutase (Gibco), resuspended in DMEM medium supplemented with 10% FBS and plated on a dish. After 30 minutes, MEF feeder cells but not SG_undiff_ attached to the dish bottom firmly. Floating SG_undiff_ were collected and plated to a plate in which mitomycin C-treated Sertoli cells had been prepared beforehand. The induced germ cells were harvested for characterization. For rescue experiments, 100 nM of miRNA mimics or mRNA siRNAs were transfected into SG_undiff_ using Lipofectamine RNAiMAX reagent ten hours prior to induction (Table S4).

### Flow cytometry analysis

The cultured SG_undiff_ were treated with 0.25% trypsin to dissociate cells. The cells were filtered through a 40-μm Nylon Cell Strainer (BD Falcon), fixed in 4% PFA for 15 minutes, and permeabilized with 0.1% Triton X-100 for 30 minutes (dispensable for the anti-KIT antibody). After blocked with 5% BSA in PBS for 30 min, the cells were incubated with primary antibodies at 37°C for 1 h and then incubated with secondary antibodies at 37°C for 1 h. The analyses were preformed using a CytoFLEX research cytometer.

### RNA sequencing analysis

Total RNAs were isolated by Trizol. Two biological replicates for SG-A were used. The RNA libraries were constructed using NEBNext Ultra Directional RNA Library Prep Kit for Illumina following manufacturer's recommendations and Oligo (dT) beads (NEB) were used to isolate poly (A) mRNAs. High-throughput sequencing was performed on a Hiseq-PE150 platform.

The correlation coefficients of gene expression levels from biological duplicates were all more than 0.95. The sequencing reads were mapped to the mouse genome (UCSC, mm9) by TopHat. The mRNA expression level was represented by fragments per kilobase of transcript sequence per millions (FPKM) calculated by Cufflinks. In addition, the differentially expressed genes were identified by Cuffdiff based on the threshold of p < 0.01 together with fold change > 3/2 or < 2/3. RefSeq mRNAs downloaded from UCSC (mm9) were used as the reference mRNAs.

### GO analysis and GSEA analysis

Gene ontology (GO) analysis was performed using DAVID bioinformatics tools (Huang da et al., 2009) and GO terms with p<0.05 were considered to be significant. GSEA analysis was performed using GSEA v4.0.1 software (Subramanian et al., 2005) with FPKM data.

### Statistical analysis

All experiments reported here were repeated at least three independent times except that RNA-seq for SG-A was performed with two biological samples. All of the values in the figures are shown as mean ± SEM unless otherwise stated. Excel 2016 or GraphPad Prism 7 was used to perform statistical analyses. For statistical analysis of differences between two groups, two-tailed unpaired Student's t tests were used. For statistical analysis in Fig. 2D, Chi-square test was used. No samples or animals were excluded from analyses. Sample size estimates were not used. Mice analyzed were litter mates and sex-matched whenever possible. Investigators were not blinded to mouse genotypes or cell genotypes during experiments.

In all figures, *, ** and *** represent p < 0.05, p < 0.01, and p < 0.001, respectively. NS (not significant) is stated if the comparison is not statistically significant (p > 0.05).

## Acknowledgements

We thank David Zarkower for antibodies. We thank Shiwen Li, Shuguang Duo, Xili Zhu, Xia Yang, Hua Qin and Qing Meng in Institute of Zoology, Chinese Academy of Sciences for their technical assistance. We thank LetPub (www.letpub.com) for its linguistic assistance during the preparation of this manuscript.

## Data Availability

The RNA-seq data set from this study has been submitted to the NCBI Gene Expression Omnibus under the accession number GSE126936.

## Author Contributions

C.H. supervised the project. C.H. and J.C. conceived and designed the study, analyzed the data and wrote the manuscript. J.C., Y.N., W.H., C.Z., D.Z., L.Y., B.J. and Y.Z. performed the experiments. C.G., X.L., M.A.H. and J.C. conducted the bioinformatics analyses. All authors commented on the manuscript.

