## Supplementary information for "MicroRNA-202 prevents precocious spermatogonial differentiation and meiotic initiation during mouse spermatogenesis"

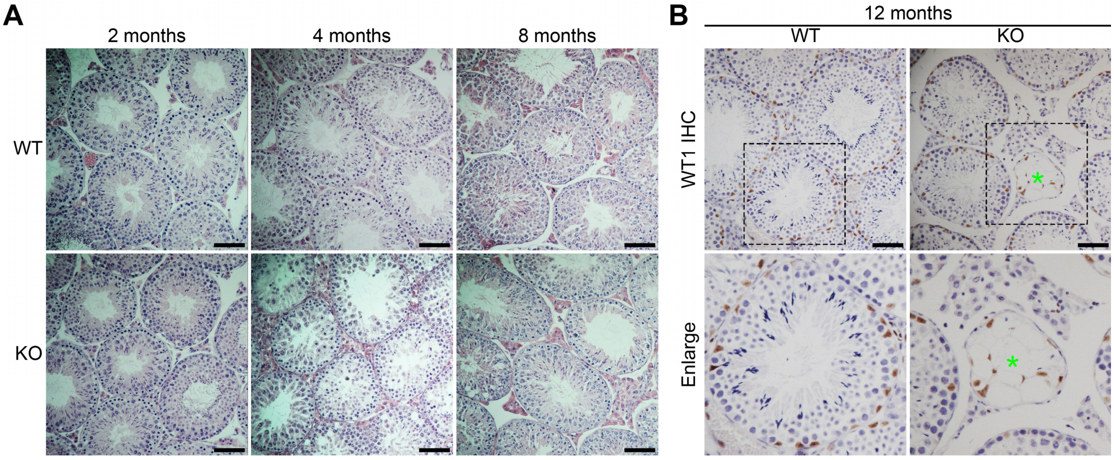


**Fig. S1.** ***miR-202*** **knockout mice show age-dependent loss of germ cells.** (**A**) Histological analysis of testis sections from WT and KO mice by H&E staining at indicated ages. (**B**) Immunohistochemistry of the Sertoli cell marker MVH in testicular sections from mice at 12 months of age. Magnified views are indicated by dashed areas. Green asterisks indicate the agametic tubules. Scale bars, 50 µm.

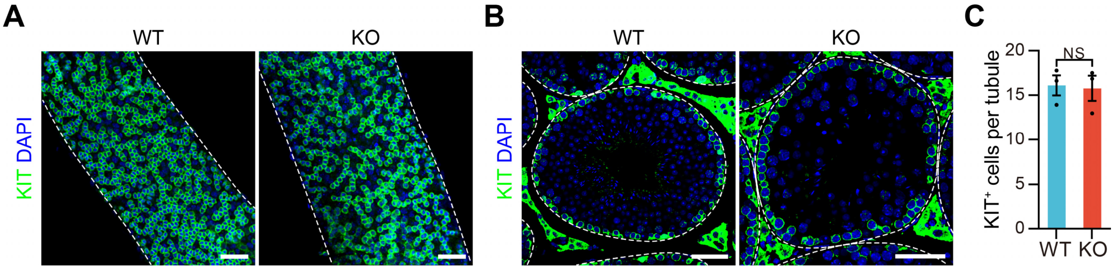


**Fig. S2.** ***miR-202*** **knockout does not disrupt spermatogonial differentiation.** (**A** and **B**) Immunofluorescent staining for KIT in whole-mount tubules (A) or sections (B) from WT and KO mice at four months of age. Dotted white line, outline of seminiferous tubules or testis tubules. (**C**) Average numbers of KIT-positive cells per tubule in (B). At least 50 tubules were counted for each mouse. All panels show mean ± SEM. NS, not significant. Scale bars, 50 µm.

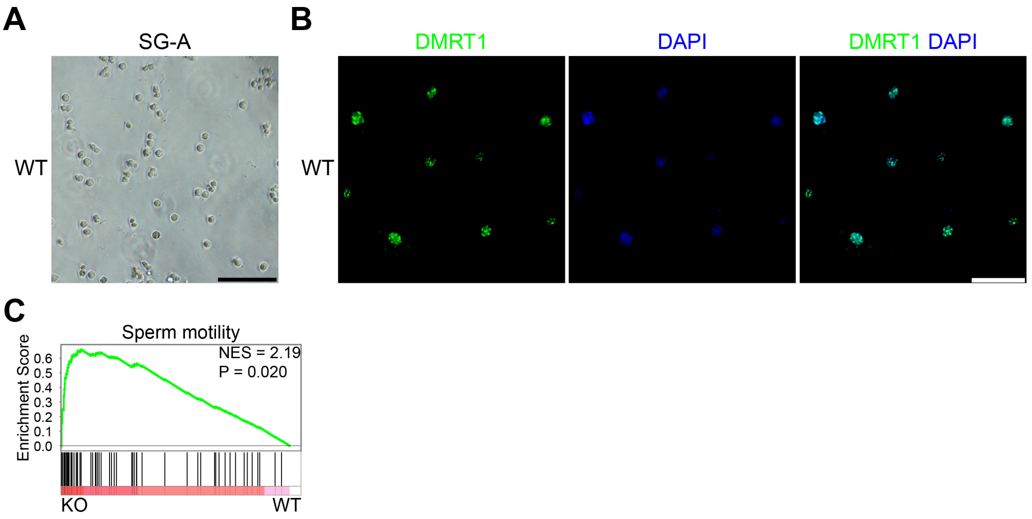


**Fig. S3. Isolation and RNA sequencing analysis of SG-A.** (**A**) Morphological evaluation of isolated SG-A by STAPUT. (**B**) Immunostaining evaluation of isolated SG-A for DMRT1. (**C**) Gene-set enrichment analyses (GSEA) for indicated GO terms in KO SG-A versus WT SG-A. On the x axis, genes are ranked on the basis of the expression ratio of KO versus WT. The nominal p value was determined by an empirical gene-set based permutation test. NES, normalized enrichment score. Scale bars, 100 μm (A) and 50 μm (B).

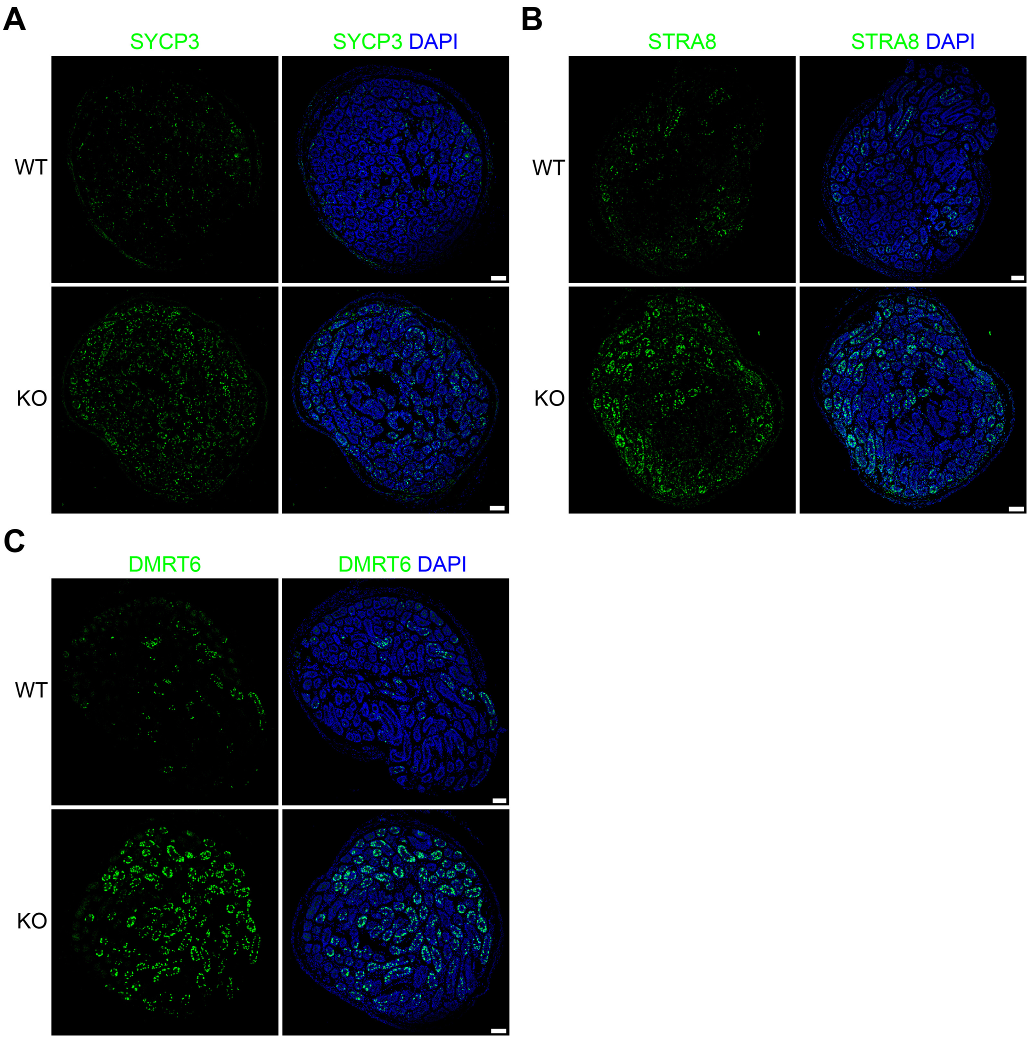


**Fig. S4. Images of whole testis sections from WT or KO mice at P9.** (**A-C**) Images of the whole sections from WT or KO mice after immunostained for SYCP3 (A), STRA8 (B) and DMRT6 (C). Note that the tubules containing SYCP3, STRA8 or DMRT6 positive cells increase significantly in KO mice. Scale bars, 100 μm.

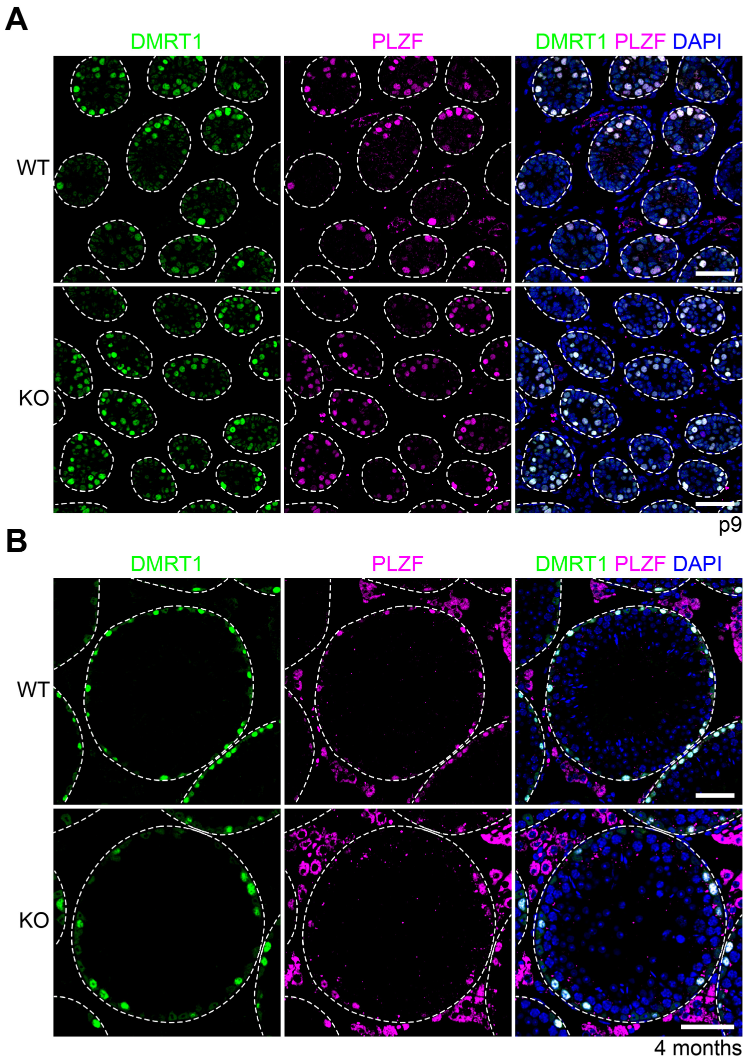


**Fig. S5. *miR-202* knockout does not affect the expression of DMRT1.** (**A** and **B**) Immunofluorescent staining for DMRT1 and PLZF in sections from mice at P9 (A) and 4 months of age (B). Dotted white line, outline of testis tubules. Scale bars, 50 µm.

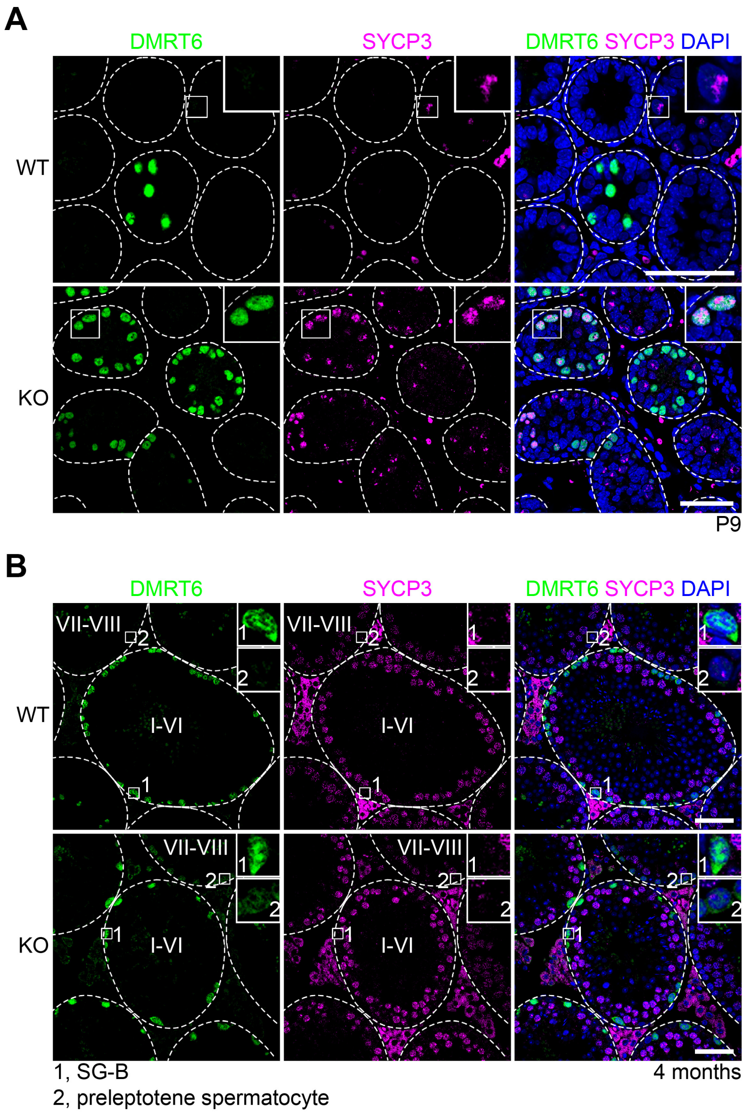


**Fig. S6. *miR-202* knockout induces the persistent expression of DMRT6 in preleptotene spermatocytes.** (**A** and **B**) Immunofluorescent staining for DMRT6 and SYCP3 in sections from mice at P9 (A) and 4 months of age (B). Insets show high magnification images of boxed regions. Dotted white line, outline of testis tubules. Scale bars, 50 μm.

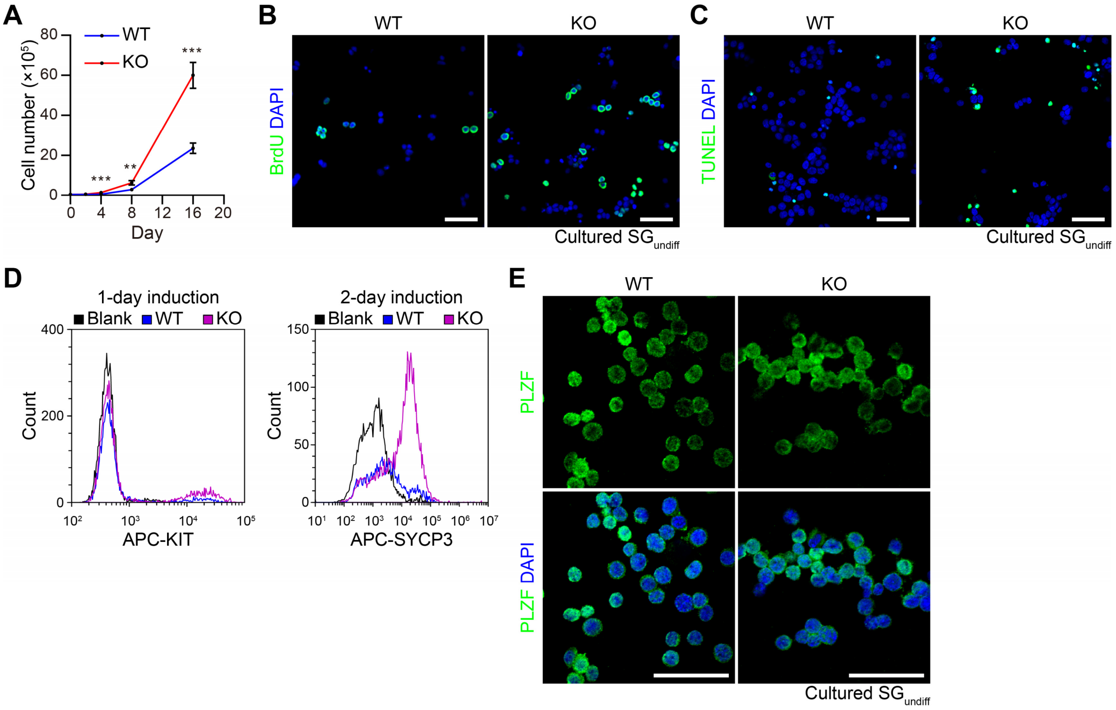


**Fig. S7. Characterization of cultured SG$_{undiff}$.** (**A**) Quantification of cell numbers for cultured WT and KO SG$_{undiff}$. n = 3. (**B** and **C**) BrdU incorporation (B) or TUNEL (C) assay for WT and KO SG$_{undiff}$. (**D**) Representative images of flow cytometry analyses for KIT and SYCP3 in WT and KO SG$_{undiff}$, after induced on Sertoli cells for one day and two days, respectively. (**E**) Immunofluorescent staining for PLZF in cultured SG$_{undiff}$. All panels show mean ± SD, $^{**}p < 0.01$, $^{***}p < 0.001$. Scale bars, 50 μm.

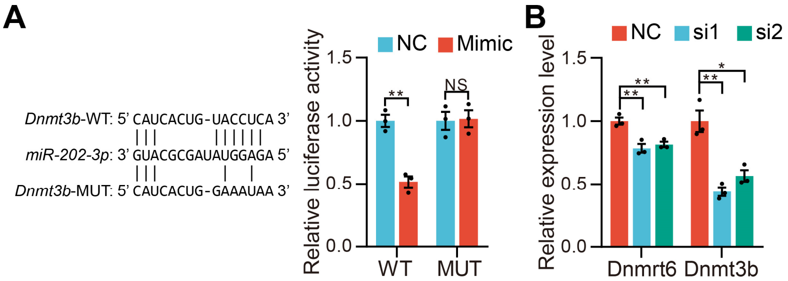


**Fig. S8. Verification of the knockdown efficiency of siRNAs for *miR-202* targets.** (**A**) Validation of *Dnmt3b* mRNA as a direct target of *miR-202-3p* by dual luciferase assay. Data are normalized to the scrambled negative control (NC) mimic. Left panel: the sequence of *miR-202-3p*, and predicted miRNA regulatory elements at the 3' UTR of *Dnmt3b* mRNA and mutated 3' UTR sequence. (**B**) qPCR analysis to verify the knockdown efficiency of siRNAs for indicated genes. Data are normalized to the NC siRNA. All panels show mean ± SEM, *p<0.05, **p < 0.01. NS, not significant.

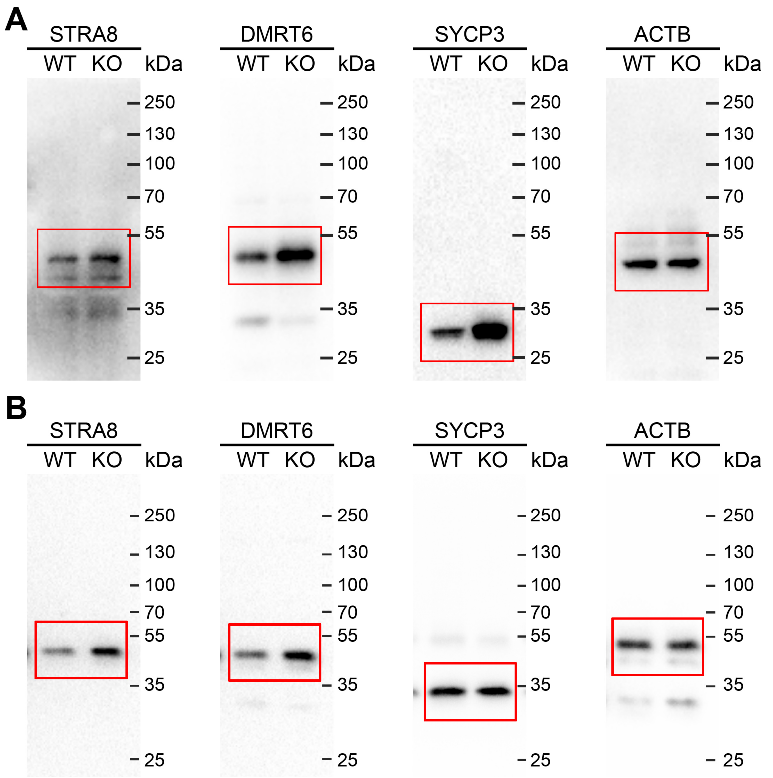


**Fig. S9. Original un-cropped Western blots shown in the manuscript.** (**A**) Western blots correspond to Fig. 2I. (**B**) Western blots correspond to Fig. 2K.

**Table S1.** RNA sequencing analysis of mRNAs in SG-A and GO terms of biological process enriched in upregulated and downregulated mRNAs in KO.

**Table S2.** List of primers.

| Target | Forward | Reverse | Application |
|---|---|---|---|
| *Actb* | CATTGCTGACAGGATGCAGAAGG | TGCTGGAAGGTGGACAGTGAGG | qRT-PCR |
| *Dazl* | AATGTTCAGTTCATGATGCTGCTC | TGTATGCTTCGGTCCACAGACT | qRT-PCR |
| *Dmc1* | CCCTCTGTGTGACAGCTCAAC | GGTCAGCAATGTCCCGAAG | qRT-PCR |
| *Dmrt6* | GAGCTTTGCTACCCGGATCA | TCCCAGCTCCTACCACATGA | qRT-PCR |
| *Dnmt3b* | CCCATGCAATGATCTCTCTAAC | AGAATGGACGGTTGTCGC | qRT-PCR |
| *Gfra1* | AGGCTCAGAATTTGTTAATGG | TAGGGCTCAAGGGAAGGAAG | qRT-PCR |
| *Id4* | CAGTGCGATATGAACGACTGC | GACTTTCTTGTTGGGCGGGAT | qRT-PCR |
| *Kit* | GCCACGTCTCAGCCATCTG | GTCGCCAGCTTCAACTATTAACT | qRT-PCR |
| *Mlh1* | CAGCTGATGGGAAATGTGCG | CAGGGTTTAGGAGGGGCTTG | qRT-PCR |
| *Mvh* | CTAGGAAGACCAAATAGTGAATCTGAC | TCCAGAACCTGTTACTACTTCTTCATT | qRT-PCR |
| *Nanos2* | CCATATGCAACTTCTGCAAGC | TGAGTGTATGAGCCTGGTCG | qRT-PCR |
| *Oct4* | GATGCTGTGAGCCAAGGCAAG | GGCTCCTGATCAACAGCATCAC | qRT-PCR |
| *Plzf* | ATTTACTGGCTCATTCAGCG | CCAGTATGGGTCTGTCTGTG | qRT-PCR |
| *Rad51* | AAGTTTTGGTCCACAGCCTATTT | CGGTGCATAAGCAACAGCC | qRT-PCR |
| *Rbfox2* | GGGATGCAGAACGAGCCAC | CTGAACCATTGCGTCAGGAG | qRT-PCR |
| *Rec8* | ATGGAGACGCTGGAAGATGC | CGGGGTTGCAGCCTCTAAA | qRT-PCR |
| *Ret* | AGGACTGGGTAGTTGCATCC | CATATATTGAGCCGAGGACAGC | qRT-PCR |
| *Sohlh1* | CGGGCCAATGAGGATTACAGA | TCCTGCGTTCTCTCTCGCT | qRT-PCR |
| *Sohlh2* | GGATTAAAGGCCCCGTTGTC | ATCGCTCTTCCTCCCCTTGA | qRT-PCR |
| *Spo11* | ACGGATCCATGTCCATGTC | TATCAGAAGGGAGGAGACCA | qRT-PCR |
| *Stra8* | ACAAGAGTGAGGCCCAGCAT | CCTCTGGATTTTCTGAGTTGCA | qRT-PCR |
| *Sycp1* | CCATGCTCGAACAGGTTGCT | ACAGTCTGCTCATTGGCTCT | qRT-PCR |

| | | | |
|---|---|---|---|
| *Sycp3* | AGCCAGTAACCAGAAAATTGAGC | CCACTGCTGCAACACATTCATA | qRT-PCR |
| *Thy1* | AACTCTTGGCACCATGAACCC | GCTGGTCACCTTCTGCCCTC | qRT-PCR |
| *Dnmt3b* | CACCTCACCTGTCCCCTTTT | ATGTGGAGCACGGGTTACTT | Amplification of mRNA 3' UTR |
| *Dmrt6* | GGCATAGATGGCGGGGGTTA | CTGTCCTGAGTGAGGCTTGG | Amplification of mRNA 3' UTR |
| *miR-202* | AAGATCCGCTTGCGTAGGAA | CCAAGCTTAGTGGGGCTCTTT | Genomic PCR |

**Table S3.** List of antibodies.

| **Antibody** | **Host** | **Company** | **Dilution** | **Cat #** |
|---|---|---|---|---|
| SYCP3 | Mouse | Abcam | IF: 1:200<br>WB: 1:2000<br>Flow: 1:100 | ab97672 |
| KIT | Goat | R&D | IF: 1:200<br>Flow: 1:100<br>Whole mount: 1:100 | AF1356 |
| MVH | Rabbit | Abcam | IHC: 1:200 | ab13840 |
| WT1 | Rabbit | Epitomics | IHC: 1:200 | 2797-1 |
| γH2AX | Rabbit | CST | IF: 1:200 | 9718 |
| PLZF | Goat | R&D | IF: 1:200<br>Whole mount: 1:100 | AF2944 |
| STRA8 | Rabbit | Abcam | IF: 1:200<br>WB: 1:2000 | ab49405 |
| BrdU | Mouse | BD Pharmingen | IF: 1:200 | 555627 |
| ACTB | Mouse | Biodragon | WB: 1:5000 | B1029 |
| DMRT6 | Rabbit | Kindly provided by David Zarkower, University of Minnesota | IF: 1:100<br>WB: 1:1000 | NA |
| DMRT1 | Rabbit | Kindly provided by David Zarkower, University of Minnesota | IF: 1:100 | NA |

**Table S4.** List of mimic and siRNAs

| Target | Sequence (5' to 3') |
|---|---|
| *miR-202-5p* mimic | UUCCUAUGCAUAUACUUCUUU |
| *Dmrt6* si1 | GCCCAGAAGGUUCUCAGAA |
| *Dmrt6* si2 | GGCAUCCUCUGAGAUUCAA |
| *Dnmt3b* si1 | GCGGGUAUGAGGAGUGCAUUA |
| *Dnmt3b* si2 | GGAUGCUAUUGUGAAUGUG |
